# Omics Discovery Index - Discovering and Linking Public ‘Omics’ Datasets

**DOI:** 10.1101/049205

**Authors:** Yasset Perez-Riverol, Mingze Bai, Felipe da Veiga Leprevost, Silvano Squizzato, Young Mi Park, Kenneth Haug, Adam J. Carroll, Dylan Spalding, Justin Paschall, Mingxun Wang, Noemi del-Toro, Tobias Ternent, Peng Zhang, Nicola Buso, Nuno Bandeira, Eric W. Deutsch, David S Campbell, Ronald C. Beavis, Reza M. Salek, Alexey I. Nesvizhskii, Susanna-Assunta Sansone, Christoph Steinbeck, Rodrigo Lopez, Juan Antonio Vizcaíno, Peipei Ping, Henning Hermjakob

**Affiliations:** European Molecular Biology Laboratory, European Bioinformatics Institute (EMBL-EBI), Wellcome Trust Genome Campus, Hinxton, Cambridge, CB10 1SD, UK.; School of Bio-information, Chongqing University of Posts and Telecommunications, 400065 Chongqing, China.; Department of Pathology, University of Michigan, Ann Arbor, Michigan, 48109, USA.; Research School of Biology, Australian National University, Canberra, 0200, Australia.; Department of Computer Science and Engineering, University of California, San Diego, 9500, La Jolla, California 92093, USA.; Commonwealth Scientific and Industrial Research Organisation, Canberra, 0200, Australia.; Institute for Systems Biology, Seattle, Washington, USA.; Biochemistry & Medical Genetics, University of Manitoba, Winnipeg, R3T 2N2, Canada.; Oxford e-Research Centre, University of Oxford, 7 Keble Road, OX1 3QG, UK.; Department of Physiology and Department of Medicine, Division of Cardiology, David Geffen School of Medicine at UCLA, 675 Charles E. Young Drive, MRL Building, Suite 1609, Los Angeles, California 90095, USA.; National Center for Protein Sciences Beijing, No. 38, Life Science Park Road, Changping District, 102206 Beijing.

**Author notes:** These authors contributed equally to this work (Yasset Perez-Riverol, Mingze Bai). Corresponding authors: Dr. Yasset Perez-Riverol, European Molecular Biology Laboratory, European Bioinformatics Institute (EMBL-EBI), Wellcome Trust Genome Campus, Hinxton, Cambridge, CB10 1SD, UK. Henning Hermjakob European Molecular Biology Laboratory, European Bioinformatics Institute (EMBL-EBI), Wellcome Trust Genome Campus, Hinxton, Cambridge, CB10 1SD, UK.

## Abstract

Biomedical data, in particular omics datasets are being generated at an unprecedented rate. This is due to the falling costs of generating experimental data, improved accuracy and better accessibility to different omics platforms such as genomics, proteomics and metabolomics^1^,^2^. As a result, the number of deposited datasets in public repositories originating from various omics approaches has increased dramatically in recent years. With strong support from scientific journals and funders, public data sharing is increasingly considered to be a good scientific practice, facilitating the confirmation of original results, increasing the reproducibility of the analyses, enabling the exploration of new or related hypotheses, and fostering the identification of potential errors, discouraging fraud^3^. This increase in public data deposition of omics results is a good starting point, but opens up a series of new challenges. For example the research community must now find more efficient ways for storing, organizing and providing access to biomedical data across platforms. These challenges range from achieving a common representation framework for the datasets and the associated metadata from different omics fields, to the availability of efficient methods, protocols and file formats for data exchange between multiple repositories. Therefore, there is a great need for development of new platforms and applications to make possible to search datasets across different omics fields, making such information accessible to the end-user. The FAIR paradigm describes a set of guiding principles to address many of these issues, and aims to make data Findable, Accessible, Interoperable and Re-usable(https://www.force11.org/group/fairgroup/fairprinciples).

---

For the first of those principles (‘Findable’), most of the available resources for the scientific community nowadays are field specific, i.e., devoted to genomics, proteomics or metabolomics experimental datasets. In contrast to searching for a publication, finding a dataset can be a troublesome process that can involve several steps, including the search in individual repositories (including several different prominent resources per field), read the related publications, or to use a generic search engine, such as Google in hope to find the right dataset. The situation has improved in recent years thanks to the development of Consortia integrating resources within the same omics field. The ProteomeXchange^4^ and MetabolomeXchange consortia are two prominent examples in the case of metabolomics and proteomics datasets. Recently *Nature*^1^ and *Nature Biotechnology*^5^ have highlighted the need for such integration frameworks for datasets where the scientific community can find the data. Additionally, in the context of the European ELIXIR (https://www.elixir-europe.org/) and USA Big Data to Knowledge (BD2K)^6^ trans-NIH initiative, the need for a dedicated platform, search engine and service enabling the aggregation of omics datasets coming from different resources across-disciplines has been recognised, analogously to resources such as PubMed^7^ or Europe PubMed Central (EuroPMC)^8^for the scientific literature.

In this context, we introduce the Omics Discovery Index (OmicsDI -http://www.ebi.ac.uk/Tools/omicsdi), an integrated and open source platform facilitating the access and dissemination of omics datasets. OmicsDI provides a unique infrastructure to integrate datasets coming from multiple omics studies, including at present proteomics, genomics and metabolomics, as a distributed resource (**Figure 1**). To date, nine resources have agreed on a common metadata structure and exchange format, and have contributed to OmicsDI (**Supplementary Note, section 1**), including (i) the major proteomics databases: the PRoteomics IDEntifications (PRIDE) database, PeptideAtlas, the Mass spectrometry Interactive Virtual Environment (MassIVE) and the Global Proteome Machine Database (GPMDB); (ii) the three major metabolomics databases: MetaboLights, the Global Natural Products Social (GNPS) Molecular Networking and the Metabolomics Workbench; and (iii) the European Genome-Phenome Archive (EGA), the major European archive for genomics and phenotypic data. In addition, it includes a University smaller-scale database (MetabolomeExpress) proving that future resources and databases can be easily included in the framework. OmicsDI stores metadata coming from the public datasets from every resource using an efficient indexing system (**Figure 1**), which is able to integrate different biological entities including genes, proteins and metabolites with the relevant life science literature (**Supplementary Note, section 4.3**). OmicsDI is updated daily, as new datasets get publicly available in the contributing repositories.

**Figure 1.**
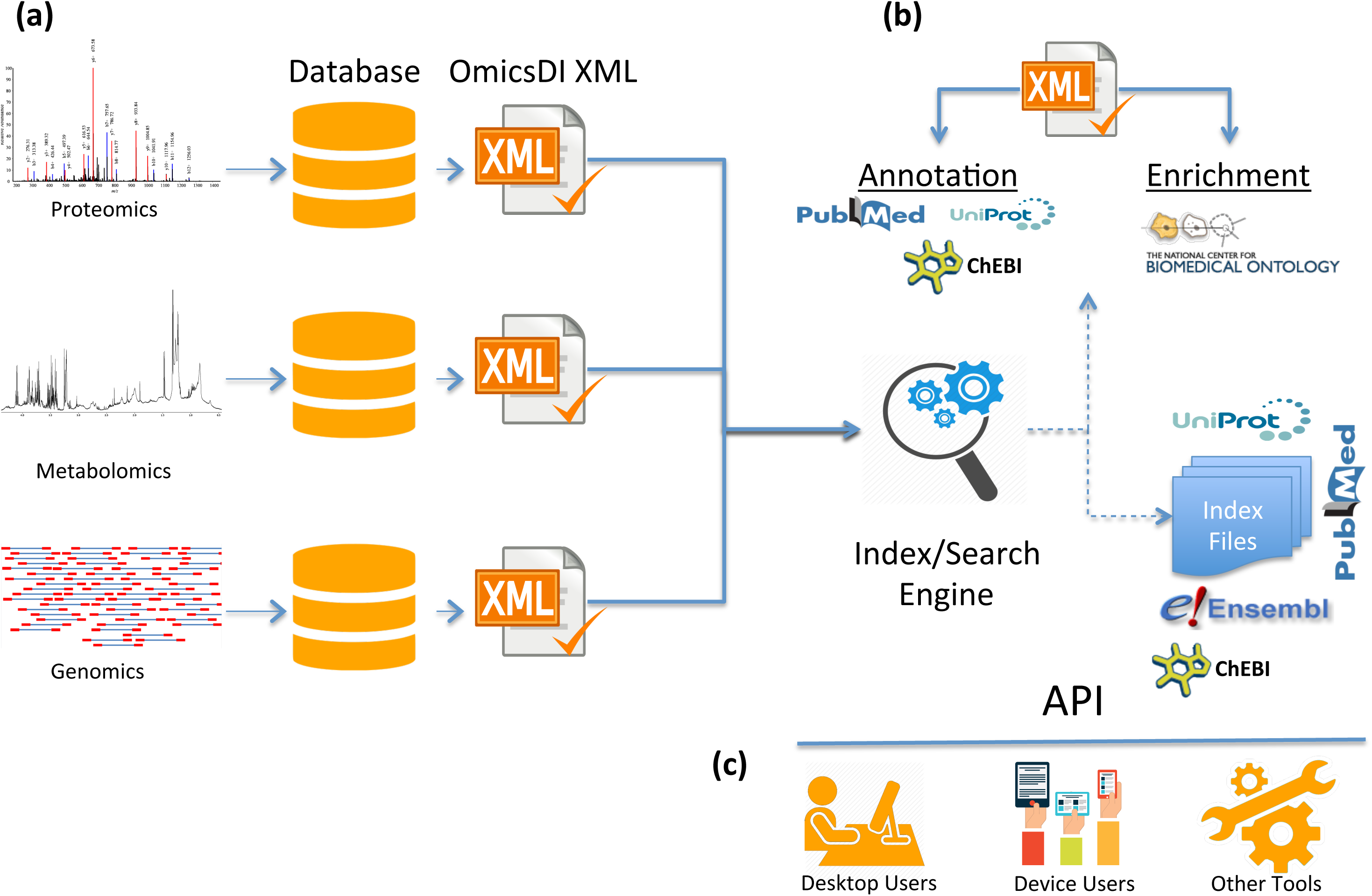
Omics Discovery Index data standardization, annotation, index and presentation: (**a**) The proteomics, genomics and metabolomics datasets stored in public repositories are converted to a common data representation including all metadata and biological entities. The OmicsDI XML files are validated using the OmicsDI XML validator; **(b)** The OmicsDI XMLs files are then annotated using public services and databases like UniProt, ChEBI or PubMed, and the metadata is enriched using the Annotator and Recommender services. The EBI Search engine generates the indexes including other related resources such as PubMed, UniProt, Ensembl or ChEBI and provides the search engine; **(c)** Different clients can use the OmicsDI API to retrieve data from the resource including the web interface and the ddiR package.

The initial list of resources represented here is just a starting point. Individual resources (including those from not yet covered omics fields) can join the OmicsDI platform by implementing the OmicsDI data integration guidelines, and metadata requirements (**Supplementary Note, section 1-2**). For this first implementation, the mandatory information per dataset includes: the data repository identifier, title, submission date, submitter information, and the URL (Uniform Resource Identifier) where the dataset can be accessed on-line. This is the minimum information that a resource needs to provide in order to make its datasets available *via* OmicsDI. In addition, two extra data types have been introduced to construct a flexible data model: ‘recommended’ metadata and ‘additional’ fields. The recommended metadata information includes at present a summary description of the dataset, sample and data protocol information, including the corresponding PubMed identifier and/or DOI (Digital Object Identifier) of the related publication, and sample annotation related to disease, tissue and taxonomy. The additional fields that can be provided are at present: protein identifiers, metabolite/small molecule identifiers, post-translational modifications, instrument platform, and annotation tags.

A flexible exchange system based on the OmicsDI XML format, using application programming interfaces (APIs) has been developed (**Supplementary Note, section 2**). Every resource willing to join the OmicsDI platform needs to generate such file format. For facilitating the process, a stand-alone open source Java tool has been developed, the ‘OmicsDI XML validator’ (**Supplementary Note, section 3**). The command line tool allows metadata error detection as well as inconsistencies in the representation of the dataset, generating a simple validation report including the different errors.

One the major challenges associated with data integration across different biomedical resources is the diversity of the associated metadata within each resource and the lack of a standard. Different resources use their own data models, metadata representation, and identifiers^9^, e.g. in the case of proteins and metabolite/small molecules, or the choice of different ontologies or controlled vocabularies (CV). To address this ‘Interoperability’ problem, OmicsDI includes a metadata normalization and enrichment step for every dataset that gets integrated into the resource. The harmonization steps standardize as much as possible the experimental and technical metadata, the identifiers for the biological entities, and the references to the external resources. For example, for any publication (DOI, citation or reference) the matching PubMed identifier has been used instead (**Supplementary Note, section 4.1**). A more difficult issue is that different datasets can have different terms representing the same ideas/concept in the same context^9^ (e.g. a protein can also be referenced as a gene product). An ontology-based enrichment step is performed using ontology tools like ‘Annotator’ and ‘Recommender’^10^, and every relevant phrase in the metadata is enriched with the relevant synonyms and ontology/CV terms(**Supplementary Note, section 4.2**). This functionality enables the users to find and associate datasets that are not even possible to find following the same approach in the original resources where they are stored. As an example, the proteomics dataset PXD001416 (http://www.ebi.ac.uk/pride/archive/projects/PXD001416), is found in OmicsDI with the search term “side effects”, but not in the PRIDE source database, which only finds the dataset with the originally annotated term “adverse effects”. In another example, it is possible to find the Metabolomics Workbench dataset ST000113 in OmicsDI using the metabolite name “Arg-[13C,15N]3”, but not in the original resource.

In addition to the use of synonyms, the OmicsDI platform expands the ‘Findable’ principle by including the ‘similar dataset’ concept. The concept of ‘Related article’ has proven to be extremely useful for other services such as PubMed to explore topic-related documents in biomedical abstracts^11^. In OmicsDI, similar datasets are computed at two different levels: metadata and biological entities (**Supplementary Note, section 5**). Both similarity levels are estimated by comparing the weighted term vectors of each dataset using the dot product. Thus, OmicsDI boosts the discoverability of related datasets (for instance datasets that use similar analytical protocols, software, or share similar biological entities), enabling the association of similar datasets stored in different resources for the very first time (**Supplementary Note, section 5**). For example, for the MetaboLights dataset MTBLS169 (http://www.ebi.ac.uk/Tools/omicsdi/#/dataset/metabolights_dataset/MTBLS169), OmicsDI reports two related datasets using the limited available metadata annotation (**Supplementary Note, section 5.1**). In contrast, the biological similarity score computes the number of shared biological entities among datasets without taking into account additional metadata. For example, the PRIDE dataset PRD000269 (http://www.ebi.ac.uk/Tools/omicsdi/#/dataset/pride/PRD000269) has a biological similarity score above 0.80 with several proteomics studies performed in aorta (**Supplementary Note, section 5.2**).

In addition to fully open datasets, life science often produces valuable datasets containing personally identifiable genetic and phenotypic data. These are managed in controlled-access repositories, such as the EGA and dbGaP, where access is granted to specified researchers after application to a data access committee (DAC). However, their metadata is accessible, and OmicsDI integrates data from EGA, as a first of such resources with open searchable metadata, but protected data. Thus, discoverability of these often very large and complex studies is increased, also through discovery by similarity from related, open datasets. This also allows OmicsDI to fully support the inclusion of increasingly common multi-omics or multi-cohort studies, where part of the data from one large collaborative effort is fully open access, while identifiable data are held under controlled-access requiring approval, e.g. Genome of the Netherlands.

Researchers and funders often need to be able to have mechanisms to track data usage over long period of time. OmicsDI tracks access to a dataset and provides a list of the most accessed datasets (**Supplementary Note, section 6.2**), providing a first solution to determine the relevance of the datasets, as previously proposed in other fields^12^. Additionally, by using cross-references between datasets, the system can track the reuse of a particular dataset across different resources. As an example, for PRIDE dataset PRD000269, OmicsDI reports both the originally deposited dataset and the reanalysed version in GPMDB.

The OmicsDI web interface provides different views, each focusing on a specific aspect of the data(**Supplementary Note, section 6.2**). A metadata overview and access statistics provide a convenient entry point to browse the resource (**Figure 2a**). Datasets can be searched and filtered based on different metadata elements such as sample attributes (e.g. species, tissue, disease),instrumentation (for example mass spectrometer type), year of publication, omics type or repository. The result of each search shows all the relevant datasets sorted using a weighted scoring function (**Supplementary Note, 6.2**). In addition, OmicsDI provides a dataset centric page, including a list of related publications and similar datasets (**Figure 2b-c**), improving the discoverability of related experimental datasets. A chord diagram presents the link between such datasets with biological similarity scores (**Figure 2c**).

**Figure 2.**
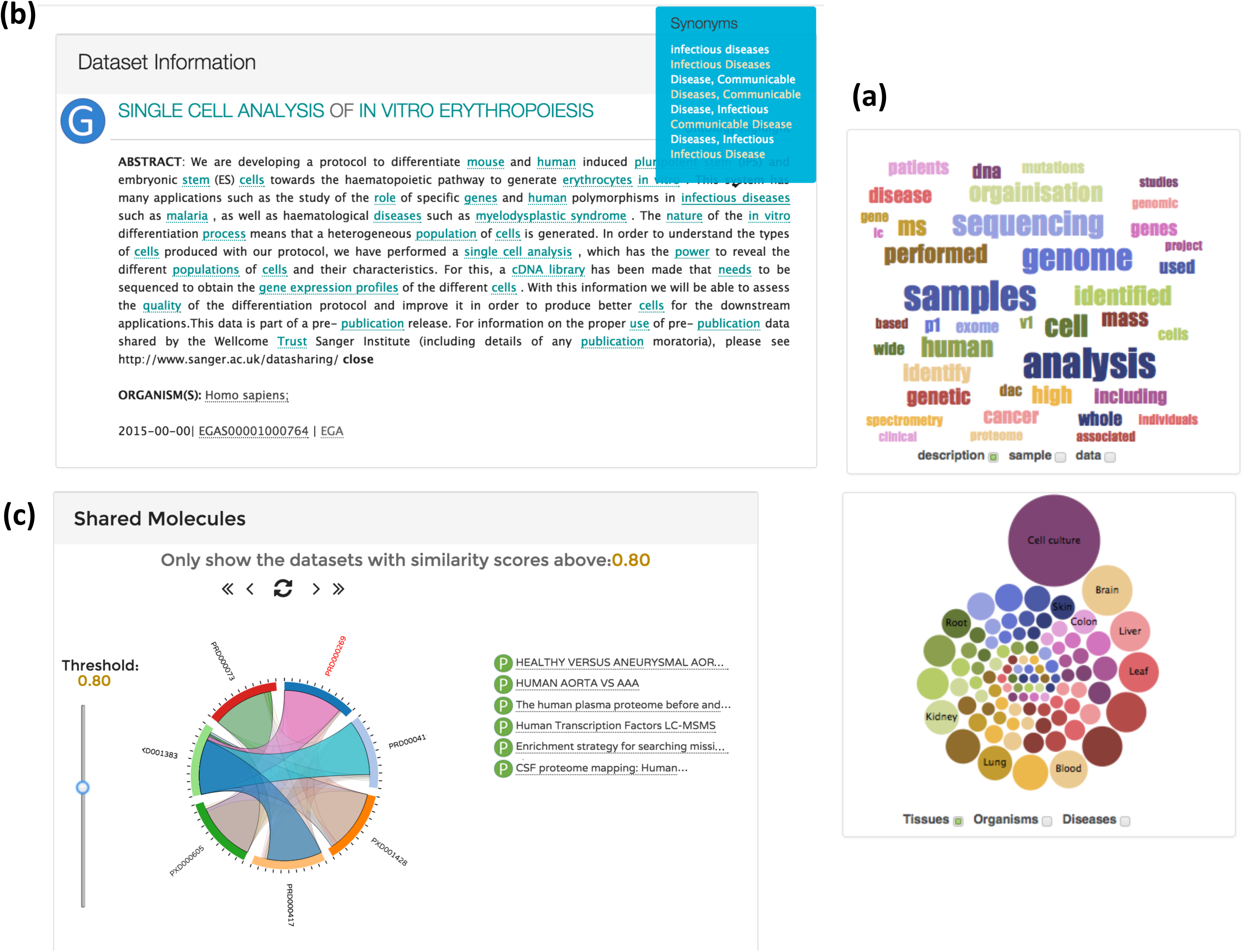
Screenshots of the OmicsDI web interface: **(a)** WordCloud and bubble plots on the home page; **(b)** The dataset view showing the related publications, including the ontology highlight option to extract the most relevant terms in the metadata; **(c)** The shared molecules box shows all datasets with a biological similarity score above 0.80.

We have also provided a web service interface, a standard RESTful API that can be used to access the data programmatically (http://www.ebi.ac.uk/Tools/omicsdi/ws). Related libraries and packages used for OmicsDI are also available at https://github.com/BD2K-DDI. For instance, an R-package called *ddiR* is provided, enabling statistical data analysis (**Supplementary Note, section 7**).

OmicsDI provides a lightweight discovery tool including more than 4,768 omics datasets from nine different repositories, three different omics types, and three continents. While advanced metadata-based browsing and indexing supports dataset findability, the lightweight approach avoids the development of redundant concepts and infrastructure. The original datasets are not replicated, but referenced. In the interest of sustainability, the responsibility for provision of a well-formatted metadata records lies with the original data providers, similarly to the concept of publisher data provision to PubMed or EuroPMC. While the central provision of a common identifier scheme across OmicsDI would be easy, and would arguably promote citability of datasets, OmicsDI displays and promotes the original dataset identifiers, to avoid creation of yet another set of identifiers, but also to ensure credit attribution to the original dataset providers. It should be noted that OmicsDI has been developed to integrate well with large, broader scope efforts like bioCADDIE (biomedical healthCAre Data Discovery andIndex Ecosystem, https://biocaddie.org/) through shared metadata formats. OmicsDI does not only provide an integrated search framework for datasets. Furthermore, as key features, OmicsDI pilots for the very first time a range of modern features in the omics domain like access metrics and discovery of related datasets that we now take for granted in the case of scientific publications.

## Acknowledgements

This work has been supported by the US NIH BD2K grant U54 GM114833, the National Natural Science Foundation of China grant [61501071], AIN is supported by
 US National Institute of Health grant [R01-GM-094231], YPR is supported by BBSRC ‘PROCESS’ grant [BB/K01997X/1]. MW is supported by NIH grant [5P41GM103484-07]. JAV and NdT are supported by the Wellcome Trust [grant WT101477MA]. TT is supported by the BBSRC ‘ProteoGenomics’grant [BB/L024225/1]. EWD and DSC. are supported in part by grant [U24 AI117966-02S1].

## Author’s contributions

HH, YPR and PP developed the OmicsDI concept. YPR and MB designed and developed the web and annotation/enrichment framework. FPV, RCB and YPR developed the GPMDB reader. SS, YMP and NB developed the indexing system based on EBI Search system. YPR, AJC, KH, DS, JP, and MW developed the Metabolomics Workbench, Metabolome Express, MetaboLights, EGA and MassIVE readers and APIs, respectively. PZ helped make the MetabolomeExpress schema OmicsDI-compatible. NT, YPR and TT developed the PRIDE reader and contributed to the web development. EWD, DSC, RMS, NB, AIN, CS, RL and JAV contributed to the design of the system. YPR, MB designed and implemented the biological similarity scoring system, and YPR performed the data analysis. YPR, JAV and HH wrote the manuscript, with contributions from all authors.

## Abbreviations

API: Application Programming Interface
bioCADDIE: biomedical healthCAre Data Discovery and Index Ecosystem
CV: Controlled Vocabulary
DAC: Data Access Committee
DOI: Digital Object Identifier
EGA: European Genome–Phenome Archive
EuroPMC: Europe PubMed Central
GNPS: Global Natural Products Social Molecular Networking
GPMDB: Global Proteome Machine Database
MassIVE: Mass spectrometry Interactive Virtual Environment
PRIDE: PRoteomics IDEntifications (PRIDE) database
OmicsDI: Omics Discovery Index
URL: Uniform Resource Identifier

